# Investigation of the isomerization of *trans*- and *cis*-cinnamic acid in Arabidopsis using stable-isotope-labeled cinnamic acid isomers

**DOI:** 10.1101/2024.05.17.594695

**Authors:** Kei Tsuzuki, Taiki Suzuki, Kotaro Nishiyama, Yoshiya Seto

**Author notes:** **Corresponding author** Yoshiya Seto, Department of Agricultural Chemistry, School of Agriculture, Meiji University, 1-1-1 Higashi-mita, Tama-ku, Kawasaki-shi, Kanagawa, 214-8571, Japan.

## Abstract

Cinnamic acid (CA) is a widely distributed metabolite in plant species and is a precursor of many important plant molecules, including lignin, flavonoids, and coumarins. CA exists as both *trans* and *cis* isomers; the *trans* isomer is more stable and common in nature. Previous reports have revealed that the *cis* isomer of CA (*cis*-CA) has auxin-like activity when exogenously applied. However, it has also been reported that *cis*-CA does not function as an auxin but affects its transport. Although these reports suggest a crucial role for *cis*-CA as an endogenous signaling molecule, its exact function and mechanism of isomerization from *trans*-CA remain unclear. Here, we report the chemical synthesis of stable-isotope-labeled *trans*- and *cis*-CA. Using these labeled compounds as internal standards, we developed a quantification method of CA using LC–MS/MS. Moreover, we monitored the endogenous conversion from *trans*-to *cis*-CA using the labeled compounds, demonstrating the UV-dependent and UV-independent CA isomerization in Arabidopsis. Additionally, we identified *cis*-CA in diverse plant species, including liverwort.

## Introduction

Cinnamic acid (CA) is a widely distributed metabolite in the plant kingdom and a precursor of many important plant molecules, including lignin, flavonoids, and coumarins. CA has a sidechain structure, which contains a carbon–carbon double bond. Therefore, CA exists as both *trans* and *cis* isomers, of which the *trans* isomer is more thermodynamically stable and common in nature. It has been reported that UV irradiation of *trans*-CA induces its isomerization to *cis*-CA. Interestingly, *cis*-CA, but not *trans*-CA, exhibits auxin-like activity when exogenously applied to Arabidopsis. At low concentrations, such as less than 1 μM, *cis*-CA promotes lateral root development in Arabidopsis and *Nicotiana benthamiana*, which leads to increased leaf weights [1,2]. However, at higher concentrations, such as 10 μM, *cis*-CA severely inhibits plant growth. The structure of *cis*-CA is partially similar to that of a major auxin, indole-3-acetic acid (IAA). However, *cis*-CA does not induce auxin receptor complex formation between the degron domain of AUX/IAA7 and TIR1 or AFB5 [2]. However, *cis*-CA inhibits polar auxin transport, similar to the well-known transport inhibitor, 1-naphthylphthalamic acid (NPA) [2].

*In planta*, CA is biosynthesized from phenylalanine by phenylalanine ammonia lyase, which produces *trans*-CA. However, it is thought that a portion of the *trans* isomer is converted into the *cis* isomer in a manner dependent on UV radiation, which is present in fluorescent light and sunlight. Indeed, in *in vitro* reactions, UV irradiation of *trans*-CA stimulates its isomerization to *cis*-CA [3,4]. However, *cis*-CA exists even when Arabidopsis is grown under LED light, which does not contain the UV portion of the spectrum [5]. Therefore, the isomerization mechanism of CA *in planta* and its triggers remain unclear. The biological activity of *cis*-CA as explained above suggests *cis*-CA may function as an endogenous plant growth regulator. To understand the endogenous role of *cis*-CA and its mechanism of isomerization, it would be effective to develop a quantitative analytical method for *trans*- and *cis*-CA. In this paper, we report the chemical synthesis of stable-isotope-labeled *trans-* and *cis*-CA. Using these labeled compounds as internal standards, we developed a quantitative method for determining endogenous levels of *cis*- and *trans*-CA by using liquid chromatography–quadruple/time-of-flight tandem mass spectrometry (LC–Q-TOF-MS). Moreover, we examined the endogenous conversion of exogenously applied labeled *trans*-CA into its *cis* isomer under fluorescent (+UV) and LED (−UV) light.

## RESULTS

### Chemical synthesis of *d*_5_-labeled *trans*- and *cis*-CA

To establish a quantitative method for determining endogenous levels of *trans*- and *cis-*CA, we synthesized stable-isotope-labeled *trans*- and *cis*-CA as the internal standard. The synthetic scheme is shown in Fig.1. Starting from commercially available *d*_5_-benzaldehyde, we first extended the sidechain by the Horner–Wadsworth–Emmons reaction to yield the ethyl ester *d*_5_-*trans*-CA [6]. We then removed the ethyl group by a simple hydrolysis reaction, which yielded *d*_5_-*trans*-CA. The MeOH solution of *d*_5_-*trans*-CA was irradiated with 254-nm UV light to obtain the *cis* isomer of *d*_5_-CA. The resulting *d*_5_-*cis*-CA was purified by HPLC. The purity of each isomer was determined to be >99% by NMR for both the *trans* and *cis* isomers (Fig. S1).

**Fig. 1.**
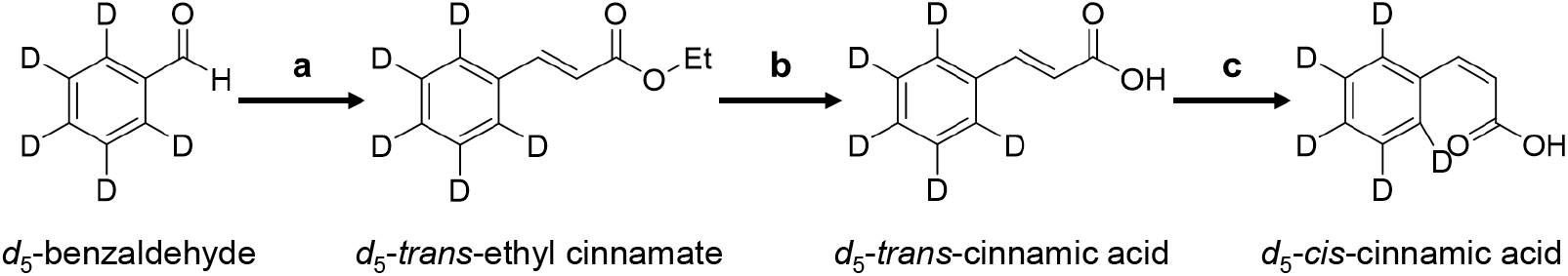
Synthesis of *d*_5_-*trans*/*cis*-cinnamic acid. (a) triethyl phosphonoacetate, NaH, THF, room temperature, quantitative; (b) EtOH, 5 M NaOH, room temperature, 79%; (c) MeOH, UV(254 nm), overnight.

### Evaluation of the stabilities of *trans*- and *cis*-CA

Generally, the *trans* form of a double-bond structure is more stable than its corresponding *cis* form. In some cases, a *cis*-configuration bond can be isomerized to the *trans* form by UV irradiation. Therefore, we considered the possibility that *cis*-CA is converted into *trans*-CA *in vitro*. To quantify the endogenous levels of small molecule compounds such as these, LC–MS/MS analysis is appropriate. To do this, it was necessary to prepare the samples, which usually includes several steps of purification using solid-phase extraction. Therefore, we analyzed the stability of *trans*- and *cis*-CA throughout the sample preparation process. As a result, we found that even *cis*-CA was stable during the sample preparation process (Fig. 2).

**Fig. 2.**
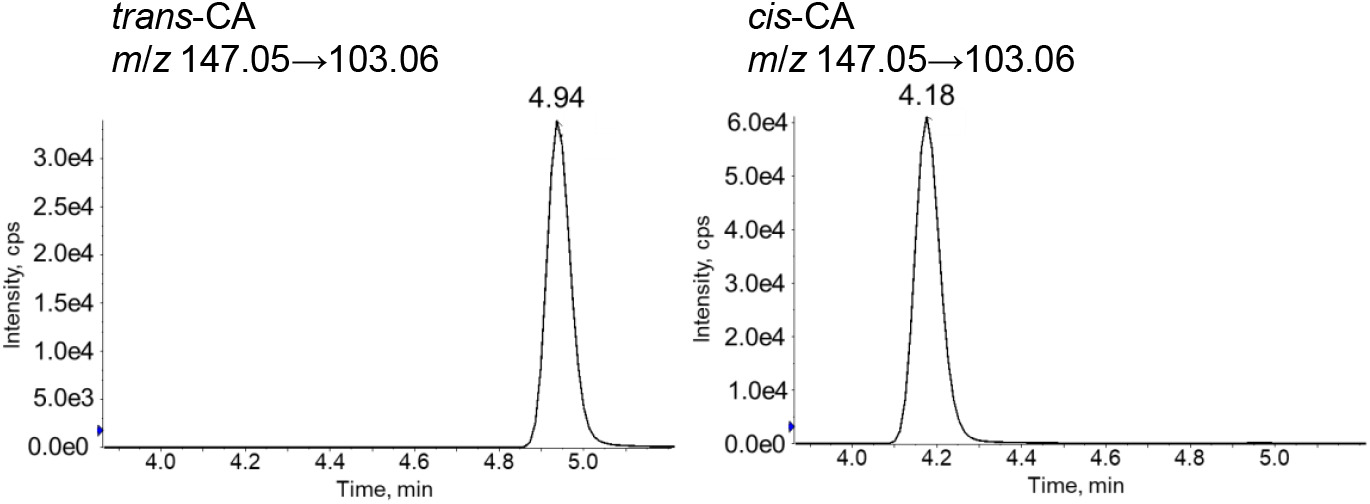
LC–MS/MS analysis of *trans*/*cis*-CA after sample preparation.

### Development of a quantitative method for determining endogenous levels of *trans-*/*cis*-CA

After confirming the stability of CA during purification, we established a quantification method for both *trans-* and *cis*-CA. In a previous report, GC was used to determine the contents of CA after methyl esterification [5]. However, it is possible that the methyl-esterified form of CA exists in plant tissue. Thus, LC–MS/MS analysis without derivatization was thought to be better for accurately measuring levels of CA. After the extraction of Arabidopsis whole plant tissue, the deuterium-labeled CA internal standards were added to the sample, and the extract was filtrated and evaporated. The samples were purified by HLB and WAX column chromatography to collect the acidic compounds, including CA. The resulting acidic fractions were analyzed by LC–MS/MS in negative-ion mode. The retention time of the *d*_5_-*trans*-CA or *d*_5_-*cis*-CA was slightly earlier than the CA peaks identified in the plant extracts because of the presence of the isotope (Fig. 3). However, the MS/MS spectrum confirmed the chemical identity of *trans*- and *cis*-CA (Fig. 3). Using this established method, we successfully detected the *cis*-CA peak in the plant extract sample (Fig. 3).

**Fig. 3.**
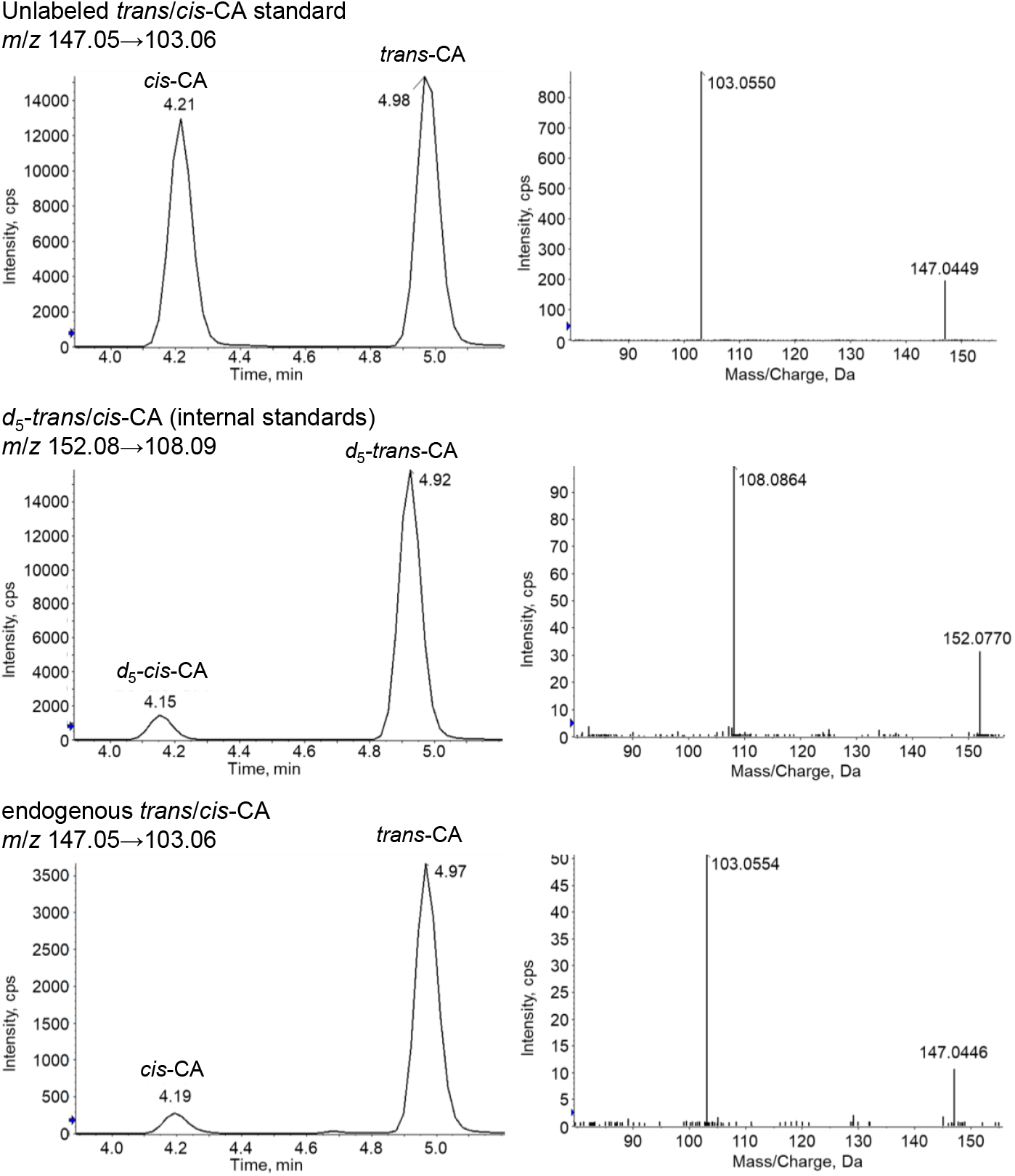
LC–MS/MS analysis of endogenous CA and *d*_5_-CA from Arabidopsis. Selected reaction monitoring (left) and full-scan spectra of fragmented ions (right) of the unlabeled authentic standard, *d*_5_-CA purified from extracts, and endogenous CA.

### Quantification of *trans*- and *cis*-CA in Arabidopsis grown under different light conditions

As mentioned above, we successfully detected endogenous *trans*- and *cis*-CA by LC–MS/MS analysis from Arabidopsis extracts. To determine whether *cis*-CA is produced only under UV-containing light, we grew Arabidopsis on MS agar medium under fluorescent light (+UV) or LED light (−UV) (Fig. S2). The shoot and root parts of two-week-old plants were used to quantify the endogenous levels of *trans*- and *cis*-CA (Fig. 4). We detected *cis*-CA in Arabidopsis grown under both fluorescent and LED light; there was no significant difference in *cis*-CA levels between the plant extracts grown under fluorescent or LED light conditions. However, by comparing the ratios of *cis*-CA in total CA, we noticed that *cis*-CA content was slightly higher when seedlings were grown under fluorescent light, and that levels of CA were much higher in roots than in shoots (Fig. 4). These results demonstrate that isomerization from *trans*-CA to *cis*-CA can occur in a UV-independent manner.

**Fig. 4.**
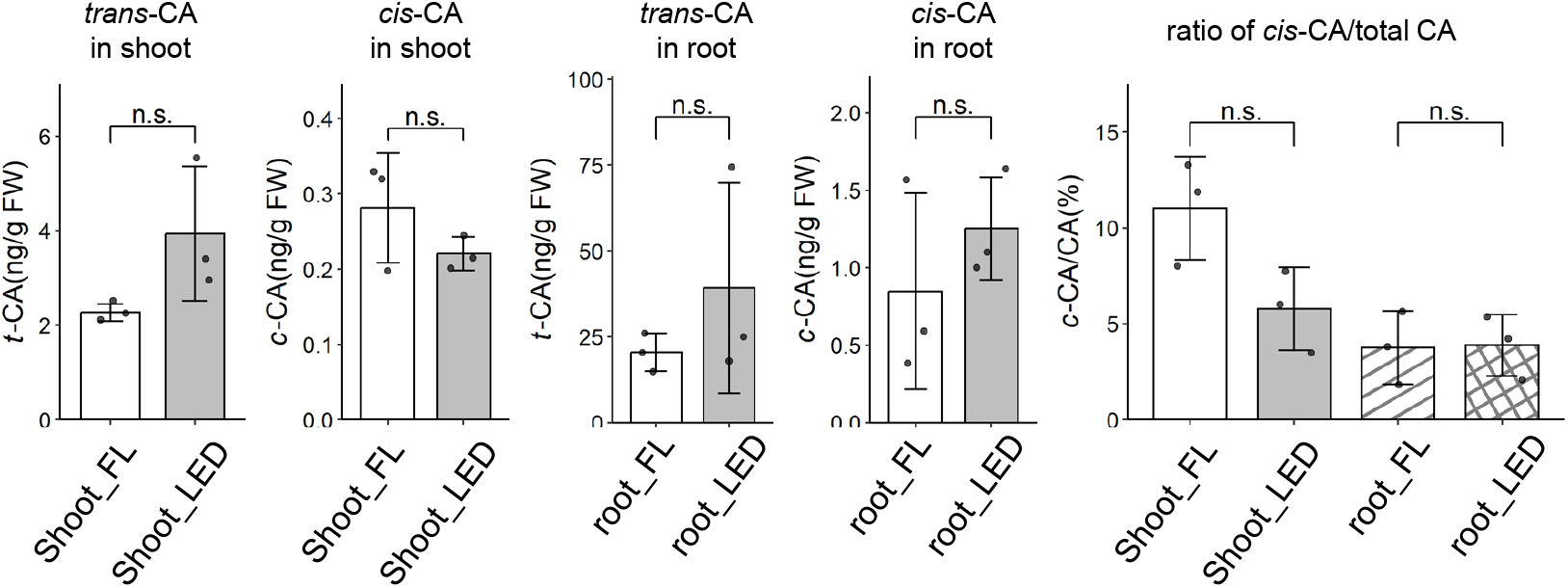
LC–MS/MS analysis of the endogenous levels of *trans*/*cis*-CA in Arabidopsis grown under fluorescent (FL) or LED light. Data are means ± SD (n=3). n.s. indicates not significant (*p*>0.05).

### Bioconversion of *trans*-CA into *cis*-CA under different light conditions

The abovementioned results suggest that the *trans*-to-*cis* isomerization of CA can occur in a UV-independent manner. To determine whether such a conversion can occur *in planta*, we performed experiments in which we fed *d*_5_-*trans*-CA to Arabidopsis Col-0 WT under either fluorescent or LED light conditions. In the negative control sample, *d*_5_-*trans*-CA incubated in water, we did not see any conversion into *cis*-CA under either light condition (Fig. 5). When *d*_5_-*trans*-CA was incubated with an Arabidopsis plant, we detected *d*_5_-*cis*-CA in both the culture medium and tissue extracts. When the plants were grown under fluorescent light, *cis*-CA was detected at much higher levels than when plants were grown under LED light (Fig. 5). These results strongly suggest that *trans*-to-*cis* isomerization of CA can occur, even under LED light conditions, but this isomerization may be accelerated by UV-light irradiation. Alternatively, it is possible there are two isomerization pathways: UV-dependent and UV-independent.

**Fig. 5.**
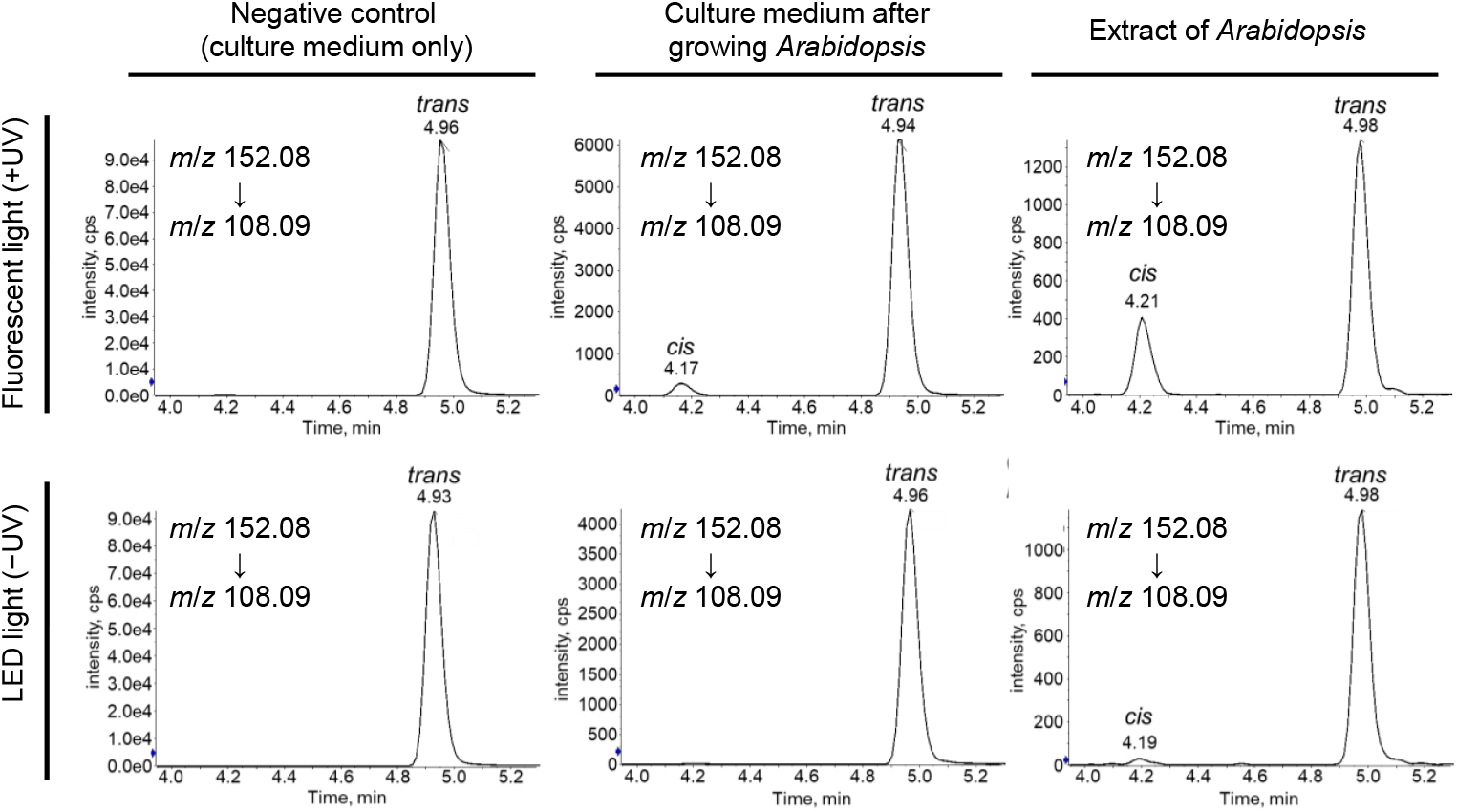
Analysis of *trans-*to*-cis* conversion of CA in Arabidopsis using *d*_5_-*trans*-CA. LC–MS/MS analysis was performed after feeding *d*_5_-*trans*-CA to Arabidopsis seedlings under either fluorescent (top three) or LED (bottom three) lights for 24 h (light/dark: 16 h/8 h).

### Distribution of *cis*- and *trans*-CA in some plant species

We then analyzed the levels of *trans*- and *cis*-CA in various plant species. We used rice (*Oryza sativa*, Nipponbare and Shiokari), tobacco (*Nicotiana benthamiana*), tomato (*Solanum lycopersicum*), and liverwort (*Marchantia polymorpha*) as plant materials. We chose both monocot and dicot, as well as a basal land plant species to see whether the CA isomerization system is conserved universally in plant kingdom. After growing for1-2 weeks, shoot and root parts were separated, and *trans-* and *cis*-CA levels were quantified using the abovementioned method. As a result, we successfully detected *cis*-CA in all the analyzed plants, including the basal land plant, *M. polymorpha* (Table 1). Interestingly, the *trans*/*cis* ratio differed among plant species. In particular, the ratio of *cis*-CA in total CA was higher in *S. lycopersicum* roots and *M. polymorpha*. Moreover, even within the same plant species, such as rice, there were significant differences in *trans*-/*cis*-CA ratios between the two cultivars tested.

**Table 1.** Quantitative analysis of CA in *O. sativa* (Nipponbare), *O. sativa* (Shiokari), *N. benthamiana, S. lycopersicum, M. polymorpha*. Data are means ± SD (n = 3).

|  |  | *trans*-CA (ng/g FW) | *cis*-CA (ng/g FW) | *cis*-CA/CA (%) |
| --- | --- | --- | --- | --- |
| *Oryza sativa* (Nipponbare) | shoot | 117.43 ± 6.38 | 0.523 ± 0.114 | 0.442 ± 0.0819 |
| | root | 15.25 ± 4.04 | 0.285 ± 0.0463 | 1.91 ± 0.472 |
| *Oryza sativa* (Shiokari) | shoot | 31.38 ± 6.06 | 0.690 ± 0.158 | 2.20 ± 0.685 |
| | root | 18.81 ± 2.00 | 0.540 ± 0.0875 | 2.81 ± 0.565 |
| *Nicotiana benthamiana* | shoot | 1.31 ± 0.209 | 0.217 ± 0.0448 | 14.2 ± 1.92 |
| | root | 7.94 ± 1.24 | 1.50 ± 0.108 | 16.0 ± 1.18 |
| *Solanum lycopersicum* | shoot | 2.50 ± 0.353 | 0.248 ± 0.0398 | 9.16 ± 2.17 |
| | root | 5.82 ± 1.56 | 2.84 ± 0.873 | 32.6 ± 1.39 |
| *Marchantia polymorpha* | | 1.24 ± 0.0971 | 0.624 ± 0.0272 | 33.4 ± 0.839 |

## DISCUSSION

In this report, we chemically synthesized deuterium-labeled *trans*- and *cis*-CA. Using these labeled compounds as internal standards, we developed a quantitative analytic method for CA using LC–MS/MS. In a previous report, CA levels were analyzed by GC–MS after methyl esterification, which cannot be distinguished from the naturally existing CA methyl ester [5]. In contrast, our method can accurately quantify the amount of endogenous *trans*- and *cis*-CA. Using this newly developed method, we simultaneously quantified the endogenous levels of *trans*-CA and *cis*-CA. The results demonstrate that levels of *cis*-CA are not increased in Arabidopsis plants grown under fluorescent light compared with plants grown under LED light. However, the feeding experiment of *d*_5_-*trans*-CA indicated that more *trans*-to-*cis* isomerization of CA occurred under fluorescent light than LED light conditions. These results suggest that CA isomerization can occur in a UV-independent manner, which may be promoted by UV irradiation. There may also be both UV-dependent and -independent mechanisms of CA isomerization *in planta*. Although further experiments are needed to understand the CA isomerization mechanism *in planta* and the exact endogenous function of *cis*-CA, our newly developed method for CA quantification could be effective for addressing these questions. Moreover, stable-isotope-labeled CAs could be an effective tool to dissect the physiological roles of CA.

Using our newly developed method, we quantified the levels of *trans*- and *cis*-CA in different plant species. Although we detected both *trans*- and *cis*-CA in all plants analyzed, we found that the ratios of *trans*/*cis*-CA differed between plant species. Between two different rice cultivars, Nipponbare contained much more *trans*-CA; thus, the *cis*-CA ratio in total CA was much lower compared with that of Shiokari. Such differences between cultivars of the same plant species may be useful for identifying the factors involved in CA isomerization.

## MATERIALS AND METHODS

### Plant materials

We used *Arabidopsis thaliana* (Col-0), rice (Nipponbare and Shiokari), tobacco (*Nicotiana benthamiana*), tomato (*Solanum lycopersicum*, moneymaker), liverwort (*Marchantia polymorpha*, TAK-1).

### Chemicals

Unlabeled *cis*-CA was chemically synthesized from *trans*-CA as we previously reported [4].

*Trans*-CA is commercially available (TCI).

### Chemical synthesis of *d*_5_-*trans*- and *cis*-CA

#### *d*_5_-*trans*-ethyl-cinnamate

THF (11 mL) and triethyl phosphonoacetate (1.4 mL, 7.06 mmol) were added to NaH (170 mg, 7.05 mmol) under an argon atmosphere and stirred on ice for 10 min. *d*5-benzaldehyde (379 mg, 3.41 mmol) in THF (5 mL) was added to the mixture, and the entire mixture was stirred at room temperature for 5 h. The reaction was quenched by adding 1 N HCl. The sample was extracted with EtOAc (3×30 mL) and washed with brine (3×90 mL). The organic layer was dried over Na_2_SO_4_ and concentrated *in vacuo*. The crude sample was purified by silica gel column chromatography (*n*-hexane/EtOAc:9/1) to obtain *d*5-*trans*-ethyl-cinnamate (quant). H NMR (300 MHz, CDCl_3_): δ 7.69 (d, *J*=16.2 Hz, 1H), 6.44 (d, *J*=15.9 Hz, 1H), 4.26 (dd, *J*=6.9, 7.2 Hz, 2H), and 1.33 (t, *J*=6.9, 3H).

#### *d*5-*trans*-CA

*d*5-*trans*-ethyl-cinnamate (657.9 mg, 3.63 mmol) was dissolved in EtOH (6.10 mL), and 5 M NaOH (6.2 mL) was added to the solution. The mixture was stirred at room temperature overnight. After the reaction, the solution was diluted to 30 mL with distilled water. The sample was extracted with EtOAc (3×30 mL), and the aqueous layer was collected. The pH of the aqueous layer was adjusted to 1 using 1 N HCl, and the sample was extracted with EtOAc (3×30 mL). The organic layer was dried over Na_2_SO_4_ and concentrated *in vacuo* to obtain *d*5-*trans*-CA (438.9 mg, 2.87 mmol, 79% yield). ^1^H NMR (300 MHz, CDCl_3_): δ 7.81 (d, *J*=16.2 Hz, 1H) and 6.46 (d, *J*=16.2 Hz, 1H). HRMS [ESI- (*m*/*z*)] calculated for (C_9_H_3_ 2H_5_O_2_ -H)^-^ 152.0765, found 152.0767.

#### *d*_5_-*cis*-CA

*d*5-*trans*-CA (438.9 mg, 2.87 mmol) was dissolved in EtOH (50 mL). The solution was placed under a UV lamp (254 nm) overnight. The solvent was evaporated *in vacuo*, and the mixture of *d*_5_-*trans*- and *cis*-CA was suspended in 10 mL of distilled water. The sample was sonicated for 5 min and then centrifuged at 18,000 g for 30 min at 4°C. The supernatant was filtered and diluted to 30 mL with distilled water. The pH was adjusted using 1 N HCl, and the sample was extracted with EtOAc (3×20 mL). The organic layer was dried over Na_2_SO_4_ and concentrated *in vacuo*. The crude sample was purified by reverse-phase HPLC (ODS SP-100, MeOH/H_2_O:6/4). A portion of the sample was purified by HPLC to give *d*5-*cis*-CA (7.1 mg, 0.046 mmol, 0.14% yield). ^1^H NMR (300 MHz, CDCl_3_): δ 7.08 (d, *J*=12.6 Hz, 1H) and 5.99 (d, *J*=12.6 Hz, 1H). HRMS [ESI- (*m*/*z*)] calculated for (C_9_H_3_ 2H_5_O_2_ -H)^-^ 152.0765, found 152.0771.

### Analysis of endogenous levels of *cis*- and *trans*-CA

Plants were grown on half-strength Murashige and Skoog medium with 1% agar for 1–2 weeks under either fluorescent light (HITACHI; FLR20S) or LED light (NK system: Plantfleck) (light/dark: 16 h/8 h). The light spectra for each light condition was measured by LA-105 (Nippon Medical & Chemical Instruments CO., Ltd, Osaka, Japan, Fig. S2). The aboveground parts and the roots were separated, and the fresh weight of each was measured. The samples were added to acetone for extraction, and the mixed solution of 100 pg/μL *d*_5_-*trans*-CA and 10 pg/μL *d*_5_-*cis*-CA were added as the internal standards. The extracts were filtered, concentrated, and dissolved in H_2_O containing 1% AcOH (1 mL). The solutions were loaded onto an HLB cartridge column (1 cc, 30 mg, Waters), washed with H_2_O containing 1% AcOH (1 mL), and eluted with 80% MeCN containing 1% AcOH (3 mL). H_2_O containing 1% AcOH (500 μL) and evaporated MeCN was added to the HLB-eluted fraction. These solutions were loaded onto a WAX cartridge column (1 cc, 30 mg, Waters), washed with H_2_O containing 1% AcOH (1 mL) and 80% MeCN (2 mL), and eluted with 80% MeCN containing 1% AcOH (2 mL). The eluates were concentrated and dissolved in 50% MeCN. The samples were subjected to LC–MS/MS to quantify the amount of endogenous *cis*- and *trans*-CA. LC–MS/MS analysis of CA and *d*_5_-CA was conducted using a quadrupole/time-of-flight tandem mass spectrometer (X500R, AB SCIEX) and an ultrahigh performance liquid chromatography (Nexera, Shimadzu) equipped with a reverse-phase column (CORTECS UPLC phenyl, 1.6 μm, *φ* 2.1×75 mm; Waters). For CA and *d*_5_-CA analysis, the elution of the samples was carried out with H_2_O containing 0.05% AcOH (solvent A) and MeCN containing 0.05% AcOH (solvent B), and the mobile phase was changed from 10% B to 40% and 100% at 5 and 7 min after the injection, respectively, at a flow rate of 0.3 mL/min. The MS/MS analysis conditions were as follows: negative-ion mode; declustering potential, – 80 V; collision energy, –15 V; and parent ions (m/z) of 147.10 for unlabeled CA and 152.10 for *d*_5_-CA.

### *d*_5_-*trans*-CA feeding experiment

*A. thaliana* plants, which were grown up for 2 weeks under either fluorescent light (+UV) or LED (–UV), were transferred to 24-well plates. Each well also contained 10 μM *d*_5_-*trans*-CA in 0.1% acetone. The plates were incubated in a growth chamber under fluorescent or LED light for 24 h (light/dark: 16 h/8 h). After the 24 h incubation, the plants were washed with sterile water and extracted with acetone. The extracts were filtered, concentrated, and H_2_O containing 1% AcOH (1 mL) was added. These samples were purified on HLB and WAX cartridge columns. The eluted WAX fraction was concentrated and dissolved in 50% MeCN. LC–MS/MS was conducted on the sample, and it was analyzed for *d*_5_-*cis*- and *trans*-CA. Purification using HLB and WAX cartridge columns and LC–MS/MS analysis of the *d*_5_-CAs were conducted using the same system as that used in the analysis of endogenous CA levels.

## Acknowledgments

This work was supported by MEXT KAKENHI (22H02276), and JST, PRESTO (JPMJPR21D9), to Y.S. We thank Candace Webb, PhD, from Edanz (https://jp.edanz.com/ac) for editing a draft of this manuscript.

