## supplemental figures for "Investigation of the isomerization of *trans*- and *cis*-cinnamic acid in Arabidopsis using stable-isotope-labeled cinnamic acid isomers"

$d_5$-*trans*-CA

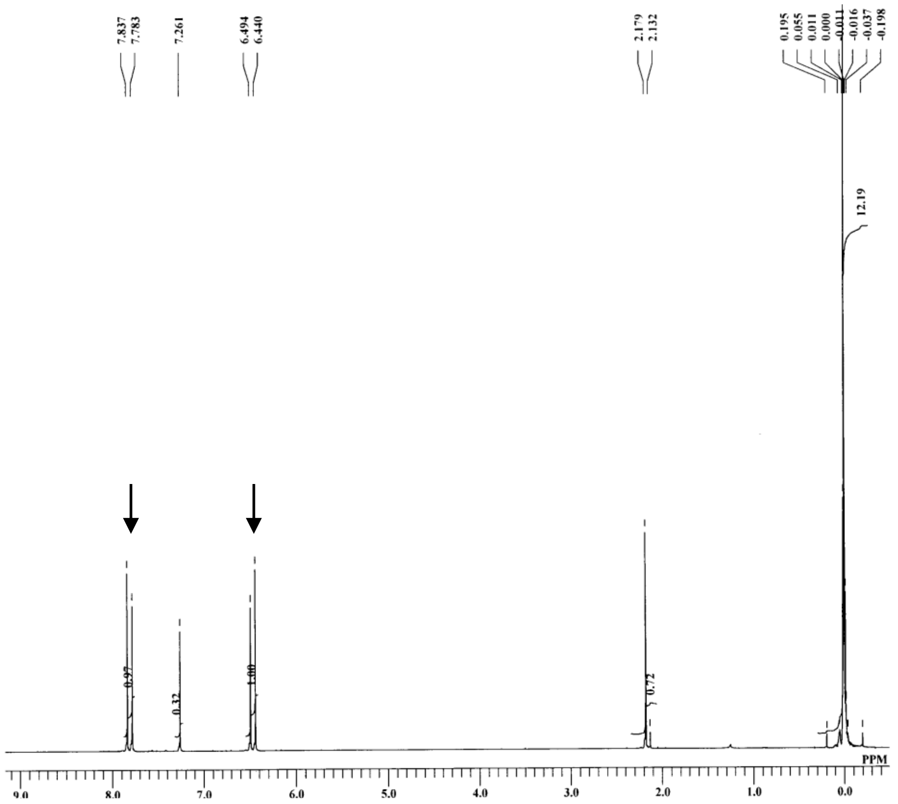

$d_5$-*cis*-CA

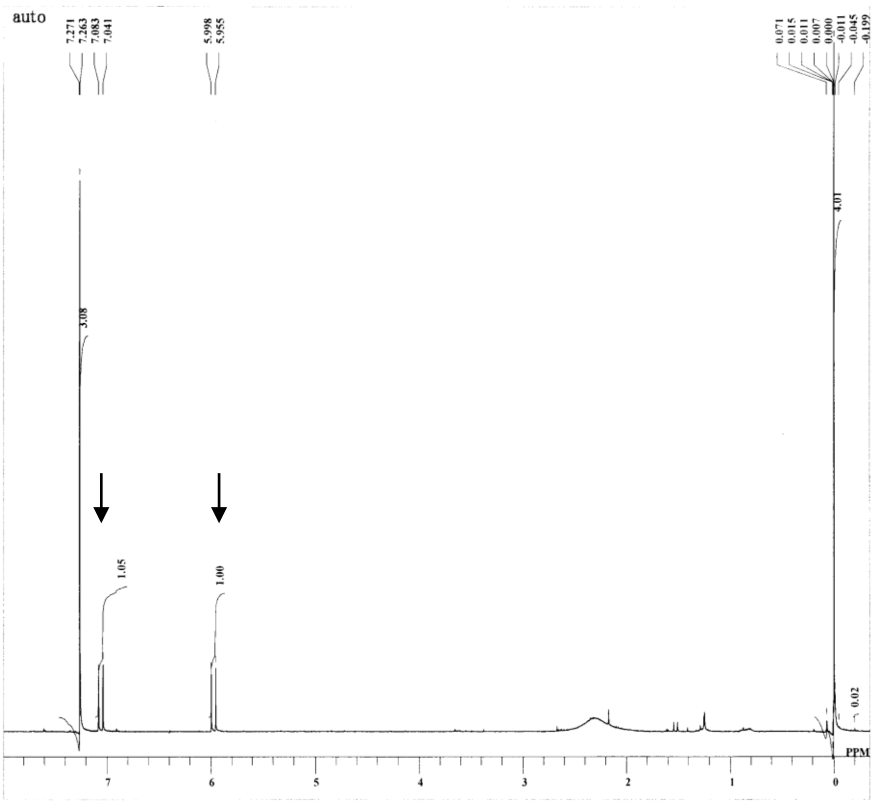

Fig.S1. [1]H-NMR analysis of deuterium-labeled *trans-*/*cis*-CA. The arrows indicate the signals derived from the protons attached on the carbon double bond in the CA side chain.

**FL**

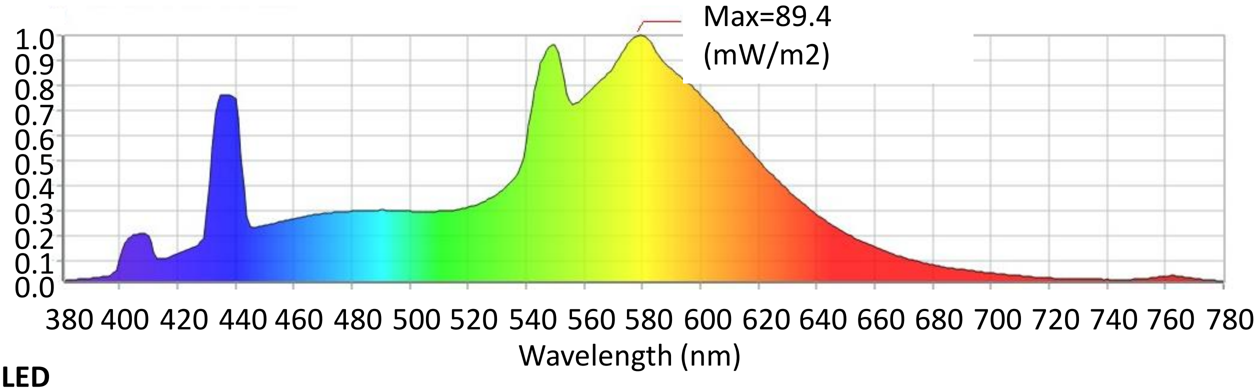


**LED**

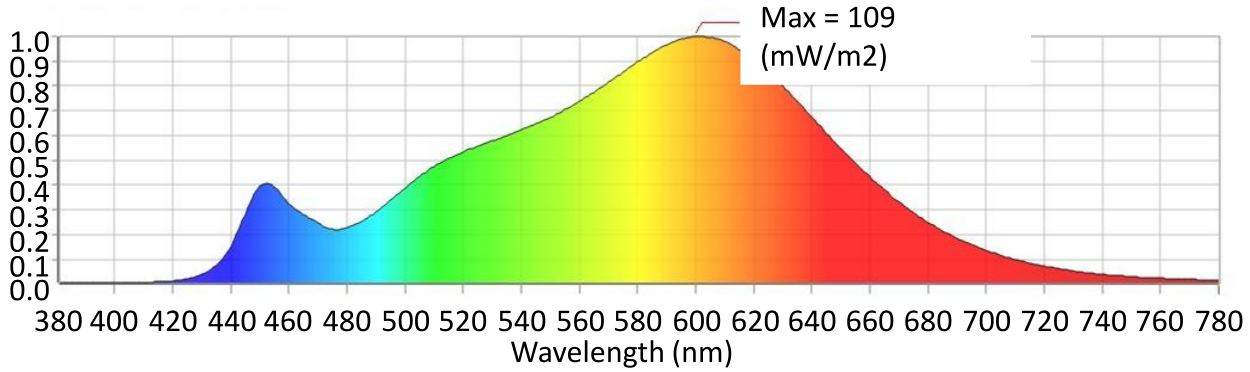


Fig. S2. The spectral photon flux density distribution of fluorescent light (FL) or LED as measured by LA-105 (Nippon Medical & Chemical Instruments Co., Ltd., Osaka, Japan).
